# Quantitative genetics of shy-bold behaviour and plastic response to novel predator cues in the cherry shrimp, *Neocaridina davidi*

**DOI:** 10.1101/2025.07.02.662724

**Authors:** Alastair J Wilson, Rosie Rickward, Francesca Santostefano

## Abstract

Understanding the genetic basis of behavioural variation among-individuals is vital for predicting if, when, and how quickly behaviour can evolve under selection. However, in heterogeneous environments, behavioural plasticity (a source of within-individual variation) may also contribute to the phenotypic variance that can be selected on. If so, a complete picture of evolutionary potential, requires estimation of genotype-by-environment interactions (GxE). Here we investigate the quantitative genetics of ‘shy-bold’ behavioural variation in the red cherry shrimp, *Neocaridina davidi,* an emerging decapod model for behavioural, genetic, and ecotoxicological research. Using a suite of behaviours associated with shy-bold personality variation we demonstrate moderate to high behavioural repeatabilities and show how a multivariate approach allows characterising the ‘shape’, not just the amount, of variation. Using a half-sib full sib breeding design in which shrimp from known families were tested under either control conditions or with predator (fish) cues present, we jointly estimate the plastic response to elevated risk, and the contribution of genetic factors to phenotypic variance. We find that genetic variance does underpin among-individual differences in behaviour. We also find evidence of plasticity, with individual shrimp shifting towards a ‘shyer’, or more risk averse, average phenotype in the presence of fish cues. However, we found no variation in plasticity either among-individuals (IxE) or among-genotypes (GxE). This implies that average behaviour can evolve under predator-mediated selection, but further adaptive evolution of behavioural plasticity may be constrained by a lack of G×E.

## Introduction

Repeatable behavioural differences among-individuals, widely referred to as personality variation (Dingemanse & Wolf, 2010a; Gosling, 2001; Réale et al., 2007; Sih et al., 2004), have been extensively documented in animal populations. This variation provides the raw material for adaptive behavioural evolution (Roche et al., 2016) in the sense that phenotypic differences among-individuals are a pre-requisite for natural selection (Houle, 1992). While there is also evidence that personality differences can have important fitness consequences in wild populations (Bell et al., 2009; Briffa & Weiss, 2010; Moiron et al., 2020; Smith & Blumstein, 2008), any response to selection depends on whether, and to what extent, personality variation is underpinned by genetic factors (Hill, 2010; Laine & van Oers, 2017). Outside of model species such as mice and zebrafish, we still know less about the genetics of animal behaviours than we do about, for example, morphology and life history traits. Nevertheless, quantitative genetic approaches have increasingly been applied to polygenic behaviours in diverse taxa, revealing that animal personality is typically heritable (Dochtermann et al., 2014). However, for plastic phenotypes expressed in heterogeneous environments, heritability may be insufficient to characterize evolutionary potential if genetic and environmental factors interact to determine phenotype rather than combining additively. This is because genotype-by-environment interactions (GxE) result in environmental sensitivity of genetic variance. Here we investigate the quantitative genetics of behavioural variation in an experimental, captive-bred population of red cherry shrimp (*Neocaridina davidi*) and ask whether experimental manipulation of (perceived) risk in the environment leads to plasticity and GxE.

The breeders’ equation of classical quantitative genetics predicts the evolution of a single polygenic trait in a constant environment as the product of a selection differential (S) and the narrow-sense heritability (Lush, 1937). Heritability (h^2^, the proportion of phenotypic variance explained by additive genetic effects; ( Falconer & Mackay, 1996) thus provides one intuitive way to characterise evolutionary potential. For behaviours, just as for any other measurable trait, additive genetic variance can be estimated using well-established statistical models, from a sample of individuals with observed phenotypes and known relatedness structure (Wilson et al., 2010). Although obtaining the high-volume data required can be logistically challenging, this approach has now been applied to many behaviours used as indicators of latent personality axes of (Réale et al., 2007). This has included work on livestock systems (e.g., aggressiveness in pigs; D’Eath et al., 2009), captive populations of wild-type animals (e.g. boldness in guppies; White & Wilson, 2019), and free-living animals studied in the wild (e.g., exploration in deer; Gervais et al., 2020).

Studies of wild populations typically report moderate behavioural heritabilities in the range of 0.2-0.5 (e.g., Brommer & Kluen, 2012; Dochtermann et al., 2014; Petelle et al., 2015). Some major empirical gaps remain. For example, genetic variation for personality has been more extensively characterised in vertebrates than in invertebrates (but see e.g., Rudin et al., 2018; Santostefano et al., 2017). Nevertheless, if personality traits are commonly under natural selection (Stirling et al., 2002), then quantitative genetic studies to date suggest genetic potential for adaptive evolution is broadly abundant.

Studies of animal personality focus on among-individual variation and, in some cases, its additive genetic component. However, since behavioural traits are highly plastic and environments are heterogeneous, we also expect phenotype to vary across observations within-individuals (Dingemanse & Wolf, 2013). Personality and plasticity are not exclusive phenomena (Dingemanse et al., 2010b; Wilson et al., 2011). For example, even consistently ‘bold’ individuals usually moderate risk-taking behaviours when environmental cues indicate heightened danger (Houslay et al., 2018). Moreover, even when exposed to identical cues individuals within a population may plastically alter their behaviour to different extents. Some individuals may be more responsive to environmental cues than others, or there could even be differences in the direction of phenotypic change. Among-individual variation in plasticity (or IxE; Nussey et al., 2007, poses a challenge to the convenient, but simplistic, idea that observed behavioural variance can be cleanly partitioned into among-and within-individual components. Similarly, where IxE is underpinned by genotype-by-environment interactions (GxE), the classic decomposition of variance into heritable (genetic) versus non-heritable (environmental) components also becomes problematic.

The evolutionary consequences of plasticity variation within populations have been explored from two major perspectives. First, in behavioural ecology, plasticity is often conceptualised as a trait that can be subject to natural selection provided it varies among-individuals (IxE; Martin et al., 2021; X Dall et al., 2019). GxE is then interpreted as heritable variation in plasticity that will facilitate a selection response. Second, and in contrast, quantitative genetic models typically propose that benefits of plasticity are selected for indirectly (sensu Lande & Arnold, 1983) as a consequence of correlations with environment-specific trait expression. For instance, if a behaviour has different phenotypic optima in different environments, then plasticity may evolve as a correlated response to environment-specific (direct) selection on the expressed trait (Gomulkiewicz & Kirkpatrick, 1992; Via & Lande, 1985). Under this second view, GxE is understood as environment-specific genetic variance. If present, GxE means the genetic variance for expressed trait (e.g. a behaviour) changes with environment and that the cross-environment genetic correlation may be <+1. In fact, the two perspectives are largely equivalent (Roff & Wilson, 2014). Thus, regardless of whether it is viewed as heritable plasticity or as environmental sensitivity of genetic variance, GxE has important consequences for adaptation in heterogeneous environments (Mulder & Bijma, 2005; Sartori et al., 2022; Toghiani et al., 2020). What remains less clear, specifically in relation to polygenic animal behaviours, is just how widespread and large GxE effects are over ecologically relevant environmental gradients.

Here we investigate the quantitative genetics of personality and plasticity in a captive population of the red cherry shrimp, *Neocaridina davidi (syn. N. heteropoda).* This freshwater Caridean shrimp is native to Taiwan but is now kept globally as an ornamental species and has become invasive in some locations via accidental or deliberate releases (Weiperth et al., 2019). The species is easy to maintain and breed under laboratory conditions and is now emerging as a decapod model for behavioural research (Azarm-Karnagh et al., 2023; Plichta et al., 2021; Takahashi, 2022a). We focus on the genetics of ‘shy-bold’ variation, an aspect of personality which describes individual propensity to engage in risky behaviour. Boldness is widely hypothesized to mediate a trade-off between resource acquisition and predation risk (Eccard et al., 2020; letters & 2007, 2007) and may also have functional importance through integration with competitive ability (Maskrey et al., 2018), stress response (Houslay et al., 2022) and/or life history strategy (Dammhahn et al., 2018). Cherry shrimp are subject to predation by ‘novel’ fish predators where populations have established outside their native range (Weiperth et al., 2019), and lab studies show increased anxiety-like (or shy) behaviours such as thigmotaxis after simulated predation attempts by net-chasing (Takahashi, 2022). While we therefore think it likely that predators mediate boldness-fitness associations in wild (or feral) cherry shrimp, our present aim is not to test hypotheses about selection, but rather to quantify the evolutionary potential of personality and plasticity. To that end we build on our earlier findings of consistent behaviour differences among individuals (Rickward et al., 2024) and characterise additive, and interactive, effects of genes and environmental factors on shy-bold behaviours. We do this by combining manipulation of predator cues with a half-sib full-sib breeding design and multivariate behavioural phenotyping. We adopt the multivariate phenotyping approach because, except in the limiting case of perfectly correlated traits, this necessarily provides more information than any single trait. In particular, by estimating the ‘shape’ of behavioural variation (White et al., 2020) we can gain deeper insights into whether, for example, among-individual variation is structurally consistent with a simple shy-bold axis (Rickward et al. 2024), or plasticity and genetic variation are directionally aligned (Berdal & Dochtermann, 2019). Here our specific objectives are to (i) test for multivariate behavioural plasticity to (perceived) predation risk and quantify IxE; (ii) estimate additive genetic variance for shy-bold personality variation; and (iii) test for and characterise the magnitude of genotype by environmental interactions (GxE) on observed behaviours in red cherry shrimp.

## Methods

### Shrimp Husbandry and Breeding

In this study we used captive-bred *Neocaridina davidi* from a colony sourced from the pet trade in February 2022 (See Rickward et al., 2024for details). To create families in a known pedigree structure, we set up 41 breeding groups, each comprising one male and four females. Except for selecting based on sex, these were sampled haphazardly sampled from stock. Each breeding group was housed in a 2.9L tank (22cm x 8.5cm x 15cm) contained within a proprietary zebrafish system (supplied by Aquaneering) with recirculating water supply. After mating, females carry the fertilised eggs attached to their swimmerets until they hatch as fully developed offspring. This meant we were able to collect full-sibling families by examining breeding groups twice weekly and transferring any ovigerous females to their own individual 2.9l tanks. Isolated females were then checked twice weekly until eggs hatched, after which the mothers were removed and retuned to stock. We recorded hatching date for family as the date hatched shrimplets were first seen. Since males remained in the breeding groups, they were able to mate with multiple females, creating paternal half-sibships. Where mortalities of males or females occurred in breeding groups, they were replaced with additional stock to maintain productivity. In total 396 adults (54 males, 342 females) were housed in breeding groups. Over a period of 8 months, we collected an offspring generation of 1191 individuals in 75 full sib-families nested within 37 paternal half-sibships. Each family was raised in its own hatching tank, containing a single plastic plant and a piece of black plastic tubing as a shelter, in one of two zebrafish systems. Within each system a recirculating water supply standardised water conditions across all tanks. Families were checked regularly and any F2 shrimplets from full-sib mating were removed before they could grow and be potentially mistaken for F1 offspring.

### Behavioural phenotyping

Behavioural phenotyping was conducted from September to December 2023, commencing when shrimp in most families had reached sufficient size for tracking (>7 mm based on prior experience). Consequently, families differed in age of phenotyping, a possible source of variation we control for statistically. Repeated observations are required to characterise among-individual variation in mean behaviour (personality) and allow for powerful within-subject tests of plasticity. However, while useful, they are not strictly required for estimation of quantitative genetic (co)variance parameters. Given the high number of shrimp in the offspring generation and some limits on researcher time and shrimp housing space, we made the pragmatic decision to generate two parallel data sets. For the first *repeated measures* data set, 80 shrimp (4 individuals haphazardly selected from each of the last 20 families produced) were assayed repeatedly. Each individual experienced six assays over a two-week period, with approximately 48 hours between assays. For the second *quantitative genetic* dataset, the remaining 1111 individuals (collectively representing all families) were each subject to a single behavioural assay.

The behavioural assay itself was identical across the two datasets. We used a modification of the standard open field test, which is widely used across animal taxa to investigate boldness and exploration and has previously been applied to cherry shrimp in Rickward et al. 2024. We added a refuge zone at one end (Figure 1), which contained artificial plants and represented 23% of the total arena floor area. Shrimp were introduced into the arena (a glass tank filled to a depth of 5cm), by being placed into a black tube positioned in the centre of the tank and allowed to acclimate for 120 seconds. The tube was then lifted out and movement tracked for a 300 s observation period. We used Sunkwang C160 video cameras mounted with a 5-50mm manual focus lens and the software Viewer II (BiObserve) to track movement while outside the refuge (note that shrimp could not be tracked in the refuge). In practice we were able to use a single camera to track in two identical arenas at once, and by duplicating this set-up were able to test four shrimp simultaneously. Arenas were surrounded by cardboard screens to prevent external visual stimuli from impacting behaviour.

**Figure 1:**
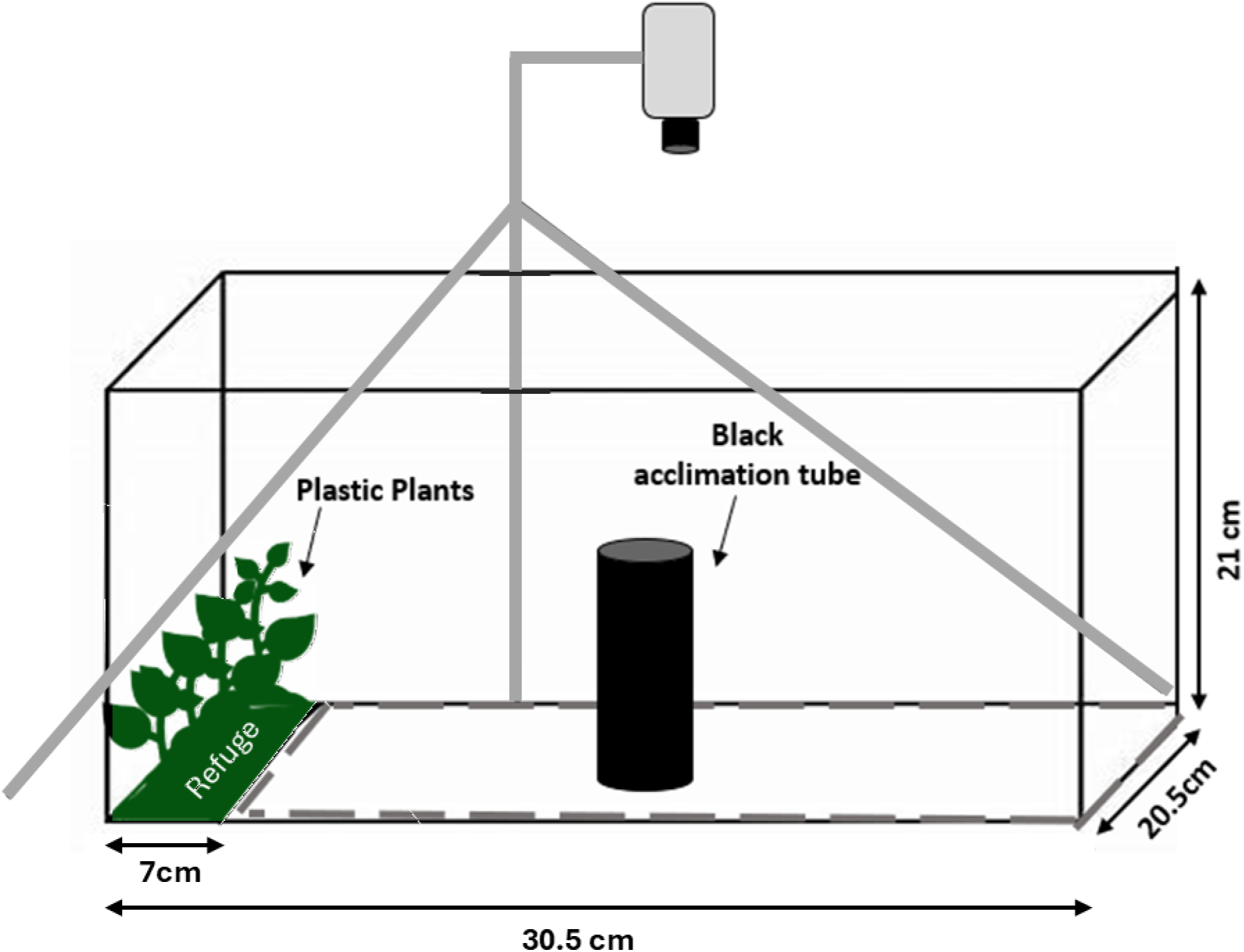
Diagram of the experimental arena used for behavioural trials.

This assay was conducted under *control* (*c*) and *predator* (*p*) treatments. In *control* conditions the arena was filled with water previously used to house conspecific shrimp. In the predator treatment we used water that had housed green swordtails (*Xiphophorus hellerii*) and so contained chemical cues of a novel predator. For the *repeated measures* dataset, each shrimp was individually housed over the testing period to allow identity to be tracked and experienced the same sequence of alternating treatment levels, starting with a control (i.e. *cpcpcp*). For the *quantitative genetic dataset* individuals remained housed in families until being tested at one treatment level only, but with *control* and *predator* assays balanced within families. At the end of each individual trial, shrimp length was measured to the nearest mm using digital callipers, and putative sex was recorded as described in Rickward et al., 2024. Rather than change water after each trial, the water was replaced after 10 consecutive trials in each arena. We also swapped the treatment level being applied in each arena between data collection sessions to ensure treatment was not confounded with any arena effects present.

We used the video tracking to extract four behavioural trait measurements from each trial; track length (cm), activity (defined as % time spent moving at >4cm.s^-1^), area covered (% of the arena explored), and duration of time spent in the refuge (s). These traits were chosen as putative indicators of shy-bold type variation (see (Rickward et al., 2024; Toms et al., 2010), with the naïve expectation that ‘bolder’ individuals will be more active and exploratory in the open arena (i.e. higher track length, activity, and area) while spending less time in the shelter. Note the traits are neither intended, nor expected, to be independent of each other. However, as described earlier, a multivariate approach provides more information and a better determination of whether the ‘shape’ of variation present is consistent with the simple shy-bold model.

### Statistical Analyses

We analysed behavioural data using univariate and multivariate (mixed) models, including pedigree-based animal models, using ASReml-R (R Core Team, 2023). We assume Gaussian residuals for all traits, an assumption that we deemed reasonable based on visual checks and after applying square root transformations to track length and activity. All (transformed) traits were scaled to standard deviation units and mean-centred on zero prior to analysis which just simplifies model fitting and results interpretation (because all four traits have the same total variance on this scaling). We denote these dependent variables as TL (track length), AC (activity), AR (area covered), and RD (refuge duration). For statistical inference on fixed effects we used conditional (Wald) F tests, while we used AIC and likelihood ratio tests (LRT) to compare among models of different random effect structure. When testing a single random effect variance using LRT, we assumed a 50:50 mix of *χ* ^2^ on 0 and 1 DF (subsequently denoted DF=0,1) following Self & Liang, 1987. For all other LRT we conservatively set the DF equal to the number of additional (co)variance parameters in the more complex of two models being compared.

### Repeated measures data set

For each behavioural trait (*y*) we fitted a series of three nested univariate linear mixed effect models with identical fixed effects but differing random effect structures. Model 1A contained no random effects, Model 1B contained a random intercept of individual identity (*id*), and Model 1C contained both a random intercept and a random ‘slope’ on treatment (specified as an interaction, *id:treatment*). Thus, the most complex model was specified as:

*y_i_ ∼ treatment + temp + order + age_family_ + n_family_ + repeat + time + arena + **id + id:treatment*** (model 1C)

Where bold font denotes random effects. Here the fixed effect of *treatment* (control vs predator) is included to test our first hypothesis that there is a within-subject plastic response to predator cues in the water. Statistical inference on *treatment* was based on Model 1C as the simpler models may pseudo-replicate this effect if either the behaviour or the *treatment* effect on behaviour varies among individuals (Schielzeth & Forstmeier, 2009). Other fixed effects were included to control for experimental sources of variation not directly relevant to our hypotheses. These included linear effects of water temperature in the arena at testing (*temp*), the *order* (1-10) of individuals tested in each arena between water changes, and the *age* at testing (which varied among families from 41 to 215 days). We also include family size (*n_family_*) at testing as a proxy of rearing density to control for any density-dependent effects on behaviour. This varied among families from 2 to 34 with a mean (SE) of 14.45 (0.392). Within-individual *repeat* number (1-6) was included to account for any systematic trend across repeated trials and *time* (of day) was also fitted. Finally, we included a fixed factor of *arena* to control for any consistent differences among the four testing arenas. Note we did not include a random *family* effect here as, beyond statistical control for age and rearing density, we wanted to estimate total among-individual variance in this dataset (as opposed to variance among individuals within families).

In models 1.B and 1.C the random intercept (*id*) partitions among-individual variance (V_I_) that can be scaled to the repeatability (R) conditional on fixed effects using R=V_I_/(V_I_+ V_R_), where V_R_ is the residual variance. Note however that under Model 1C, we denoted the two treatment levels numerically in the random part of the model, with control=0 and predator=1. This means that the intercept variance is properly interpreted as among-individual variance in the control treatment, and we thus derive an estimate of R in the control treatment.

Next, we fitted a multivariate formulation of the repeated measures mixed model to estimate the among-individual behavioural covariance matrix (**ID**). Fixed effects on each trait were as described above for the univariate case, but after finding no support for IxE in univariate models (see Results) we included random individual (*id*) intercepts only (i.e. used a multivariate formulation of 1B). This yielded an estimate of **ID** as a 4×4 matrix containing trait specific estimates of *V*_I_ on the diagonal, and estimates of COV_I_, the among-individual covariance, for each pair of traits. We standardised covariances to among-individual correlations (r_I_) to aid interpretation. Reiterating that strong among-trait correlations are expected here, we did not formally test each pairwise relationship against a null model of r_I_=0. However, we use estimated standard errors (SE) to calculate approximate 95% CI (as r_I_ ±1.96SE) to guide our interpretation. We also subjected our estimate of **ID** to eigen decomposition to summarise its structure, predicting that the first eigen vector (or principal component) of **ID,** denoted **id_max_,** will capture most of the among-individual variance, and load on all traits with signs that are antagonistic between RD (where high values indicate shyness) and the other three traits (where high values indicate boldness). We generated approximate 95% CI on eigen values trait loadings using a parametric bootstrap (Boulton et al., 2015).

### Quantitative genetic data set

We first plotted raw behavioural data by family to see if differences among-families are visually apparent before fitting a series of univariate mixed models (Models 2A – 2D) to each trait. As for the repeated measures data, these models shared a common fixed effect structure (that included *treatment, temp, order, age_family_, n_family_, time* and *arena* as defined above) with sequentially more complex random effect structures. Model 2A had no random effects, while model 2B included a random intercept of *family*. We then expanded to an animal model formulation (Wilson et al., 2010) using the known pedigree structure and adding a random genetic intercept (model 2C). A random genetic slope on *treatment* was then also added to model GxE (Model 2D). Thus, the most complex model (Model 2D) was specified for each trait (y) observed once on each individual (i) as:

*y_i_ ∼ treatment + temp + order + age_family_ + n_family_ + time + arena +* **family + a_int.i_** + **a_slp.i_**:**treatment** (Model 2D)

In this model, *a_int_* represents individual genetic merit for behaviour in control conditions. Variance in intercepts is thus interpreted as additive genetic variance in the control treatment (V_A*c*_). Similarly, a_slp_ represents genetic merit for the slope of a linear reaction norm describing the plastic response to predator cues (i.e. moving from treatment *c* to *p*). Variance in genetic slopes is thus interpreted as GxE.

Finally, we extended to the multivariate case to estimate the among trait genetic variance-covariance matrix **G**. Given little support for GxE from univariate models (see Results), in practice we fitted a multivariate version of model 2C (i.e. with genetic intercepts but not slopes). **G** includes estimates of V_A_ on the diagonal and the additive genetic covariance COV_A_ between each pair of traits off the diagonal. We rescaled covariances to genetic correlations (r_G_) to aid interpretation, and subjected **G** to eigen decomposition, facilitating comparison of its structure and shape to that of **ID**. We calculated the angle between the ***id_max_*** and ***g_max_*** (the first eigen vector of **G**), noting that this could range from 0° if the two vectors are perfectly aligned in orientation up to 90° if they are completely orthogonal.

## Results

### Repeated measures data set: Personality, plasticity, and IxE

Analysis of repeated measures data provided strong statistical support for among-individual variance in all four behaviours. For all traits, Model 1B was preferred based on AIC and a significantly better fit to the data than model 1A based on LRT (Table 1). However, inclusion of random slope variance (Model 1C) was not supported for any trait (Table 1). Indeed, slope variances (not shown) were bound to zero under standard modelling conditions in which estimates are constrained to positive parameter space. Although IxE is not supported, we nonetheless report fixed effects from Model 1C to protect against pseudoreplication. We found significant behavioural plasticity in three of the four traits. The estimated *treatment* effects on each behaviour denote the mean plastic for a transition from *control* to *predator* treatment. Thus, in combination the four *treatment* effect sizes estimated from these univariate models define the vector of average (within-individual) plasticity for the multivariate phenotype. We present this graphically in Figure 2a to facilitate visual comparison to the shape of variation revealed by multivariate models (a point explained further below). Overall, the addition of predator cues can be viewed as shifting shrimp towards a ‘shyer’ average behavioural phenotype, with mean (within-individual) decreases of approximately 0.25 standard deviation units (SDU) for both TL and AC, and a corresponding increase of 0.38 SDU for RD (Figure 2a). Note however that a more modest, and non-significant decline, is seen in AR (Figure 2a). Additional fixed effects included in these models are not directly relevant to our hypotheses and so not discussed in full here (but see supplemental table S1 for a full presentation of all effect size estimates and statistical inference).

**Figure 2:**
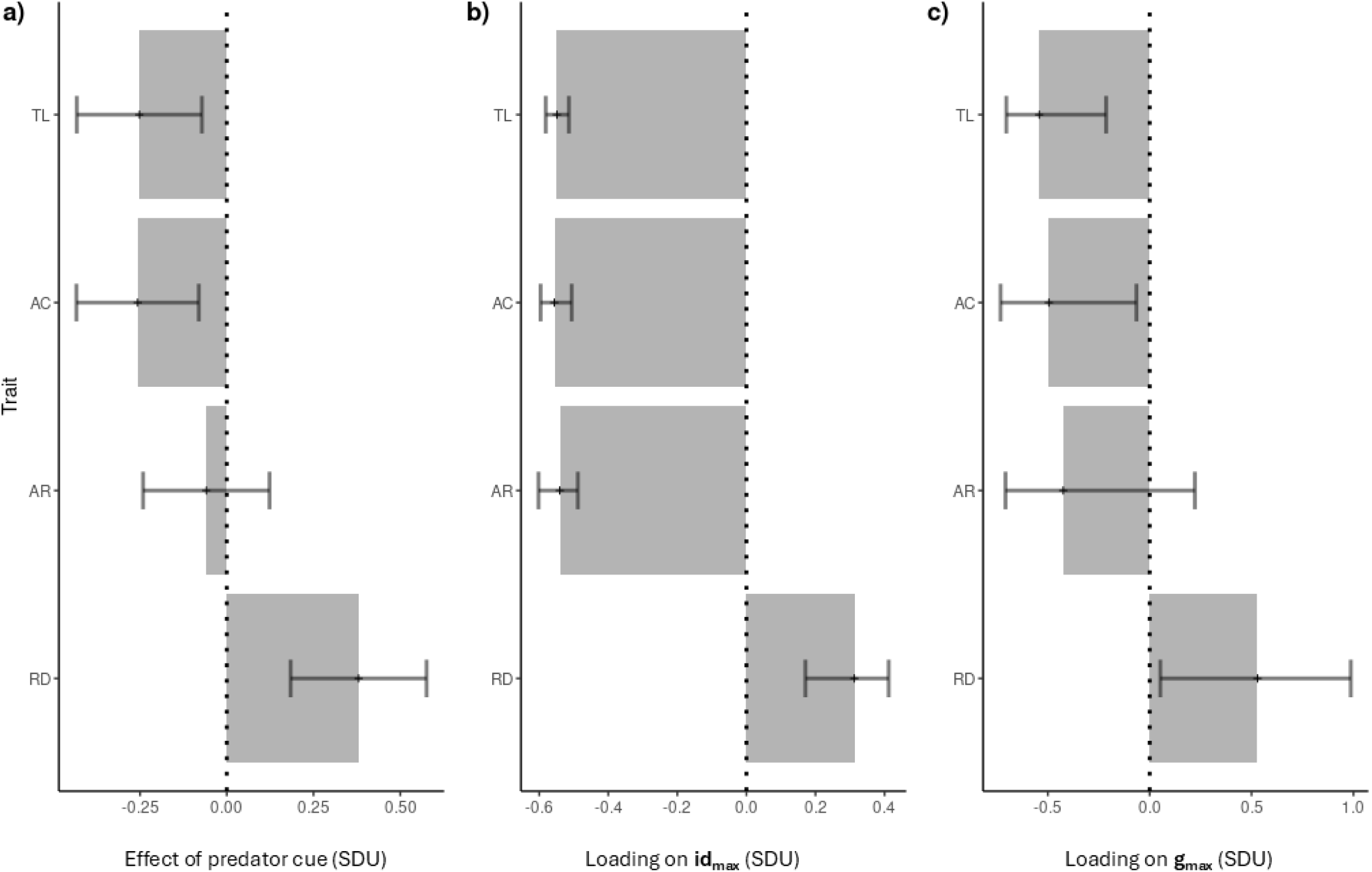
The estimated effects of predator cue presence on each trait individually are combined to define a ‘vector of plasticity’ that describes the direction and magnitude of plasticity in multivariate trait space. This vector of plasticity is represented graphically (a) alongside the leading eigen vectors of among-individual variation **id_max_** (b), and the leading vector of additive genetic variance **g_max_** (c). The vector of plasticity is estimated by extracting treatment effects from the univariate models, while **id_max_** and **g_max_** are obtained from multivariate modelling. Error bars denote 95% CI.

**Table 1:** Univariate model comparisons for analysis of *repeated measures* data showing AIC and log-likelihood (LnL) for each model fitted to each trait as well as likelihood ratio tests (LRT) of among-individual variance (V_I_) and IxE. Shading denotes preferred model (lowest AIC) for each trait.

| Trait | Model | AIC | LnL | LRT Comparisons |  |  |  |  |
| --- | --- | --- | --- | --- | --- | --- | --- | --- |
| | | | | Comparison | Tests | $\chi^2$ | DF | P |
| TL | 1A | 497.2 | -247.6 |  |  |  |  |  |
| | 1B | 425.6 | -210.8 | 1.2 vs 1.1 | $V_I$ | 73.6 | 0,1 | <0.001 |
| | 1C | 429.6 | -210.8 | 1.3 vs 1.2 | $I \times E$ | 0.010 | 2 | 0.995 |
| AC | 1A | 494.2 | -246.1 |  |  |  |  |  |
| | 1B | 415.4 | -205.7 | 1.2 vs 1.1 | $V_I$ | 80.9 | 0,1 | <0.001 |
| | 1C | 419.4 | -205.7 | 1.3 vs 1.2 | $I \times E$ | 0.001 | 2 | 1.00 |
| AR | 1A | 512.0 | -255.0 |  |  |  |  |  |
| | 1B | 440.9 | -218.5 | 1.2 vs 1.1 | $V_I$ | 73.1 | 0,1 | <0.001 |
| | 1C | 444.9 | -218.5 | 1.3 vs 1.2 | $I \times E$ | <0.001 | 2 | 1.00 |
| RD | 1A | 486.0 | -242.0 |  |  |  |  |  |
| | 1B | 464.5 | -230.3 | 1.2 vs 1.1 | $V_I$ | 23.5 | 0,1 | <0.001 |
| | 1C | 468.5 | -230.3 | 1.3 vs 1.2 | $I \times E$ | 0.005 | 2 | 0.998 |

In the absence of IxE on any trait, we used a multivariate formulation of model 1.2 (i.e. including random intercepts only). Our estimate of **ID,** which is therefore assumed to be homogenous across treatments, contains V_I_ estimates that scale to repeatabilities (SE) ranging from R = 0.174 (0.046) for RD to R=0.342 (0.053) for AC (Table 2). These are almost identical to estimates extracted from univariate models (not shown). Expressing the trace of **ID** (i.e. sum of diagonal elements) as a proportion of total variance in multivariate phenotype yields a multivariate analogue of repeatability of 0.292 (0.045). In other words, after controlling for fixed effects, individual identity explains approximately 29% of the phenotypic variance in multi-trait space. As expected, **ID** also contains strong among-trait correlation structure (Table 2), with uniformly positive corelations among TL, AC and AR which – in turn-are all negatively correlated with RD. This is reflected in the eigen decomposition which reveals that ***id_max_*** accounts for 90% of the variance in **ID** (95% CI: 84-95%). This vector loads significantly on all four original traits, with signs that are antagonistic between RD and the other traits (Figure 2b). This is consistent with a simple shy-bold paradigm in which, for example, a shrimp that has a consistently low track length in the assay, also shows consistently low activity and area covered, and spends consistently more time than average in the refuge. The similarity of ***id_max_*** (Figure 2b) to the vector of plasticity (Figure 2a), is consistent with the idea that plasticity to predator cues causes within-individual behavioural change in a direction that is closely aligned to the main axis of among-individual variation.

**Table 2:** Among-individual variance-covariance matrix **ID** estimated from repeated measures data with trait-specific V_I_ on the diagonal (black shading), among-individual covariance below the diagonal, and the corresponding correlations (r_I_) above the diagonal (grey shading). Also shown are repeatability estimates (R) conditional on fixed effects. All estimates are from a multivariate formulation of model 1.2 which assumes an absence of IxE (so that **ID** is homogenous across treatments and R_C_=R_P_). Standard errors are shown in parentheses.

| Trait | ID matrix |  |  |  | R |
| --- | --- | --- | --- | --- | --- |
|  | TL | AC | AR | RD |  |
| TL | 0.293 (0.064) | 0.991 (0.004) | 0.923 (0.026) | -0.736 (0.088) | 0.323 (0.052) |
| AC | 0.297 (0.065) | 0.308 (0.066) | 0.910 (0.027) | -0.676 (0.101) | 0.342 (0.053) |
| AR | 0.274 (0.062) | 0.277 (0.063) | 0.301 (0.066) | -0.705 (0.099) | 0.324 (0.052) |
| RD | -0.156 (0.047) | -0.147 (0.046) | -0.152 (0.046) | 0.154 (0.046) | 0.174 (0.046) |

### Quantitative genetic data set: Family effects, genetic (co)variance and GxE

Visualising the raw data revealed high variation among families in all four traits overall, while family means are moderately to strongly positively correlated across treatments (Figure 3). Accepting family differences as a first indicator of genetic variation, these visual patterns point to genetic variation in the traits but limited GxE at most (i.e. any GxE is insufficient to strongly disrupt ranking of families across treatments). More formally, univariate mixed models provide strong statistical evidence of among-family variance with Model 2B providing a significantly better fit than Model 2A in all traits (LRT, all P<0.001; Table 3). Evidence for additive genetic variance underpinning these family differences was also found, though this is slightly more equivocal. Specifically, Model 2C (animal model) was preferred based on AIC for TL, AC and RD, but the LRT comparison of models 2C and 2B was only significant at α=0.05 for TL (with marginally non-significant results for AC and RD). For AR, model 2B had the lowest AIC. More straightforwardly, and perhaps unsurprisingly given the absence of detectable IxE in the repeated measures data, we found no evidence for GxE under Model 2D. As noted previously, GxE can be modelled as genetic variance in reaction norm slope (as in Model 2D), or as environmental sensitivity of genetic variance for the trait modelled using a character state approach (Roff & Wilson, 2014). We therefore fitted additional character state models that corroborated the absence of GxE (see Supplemental Appendix), yielding treatment-specific estimates of V_A_ and **G** that very similar, and estimates of the cross-treatment genetic correlations close to +1 for four traits (Supplemental Appendix).

**Figure 3:**
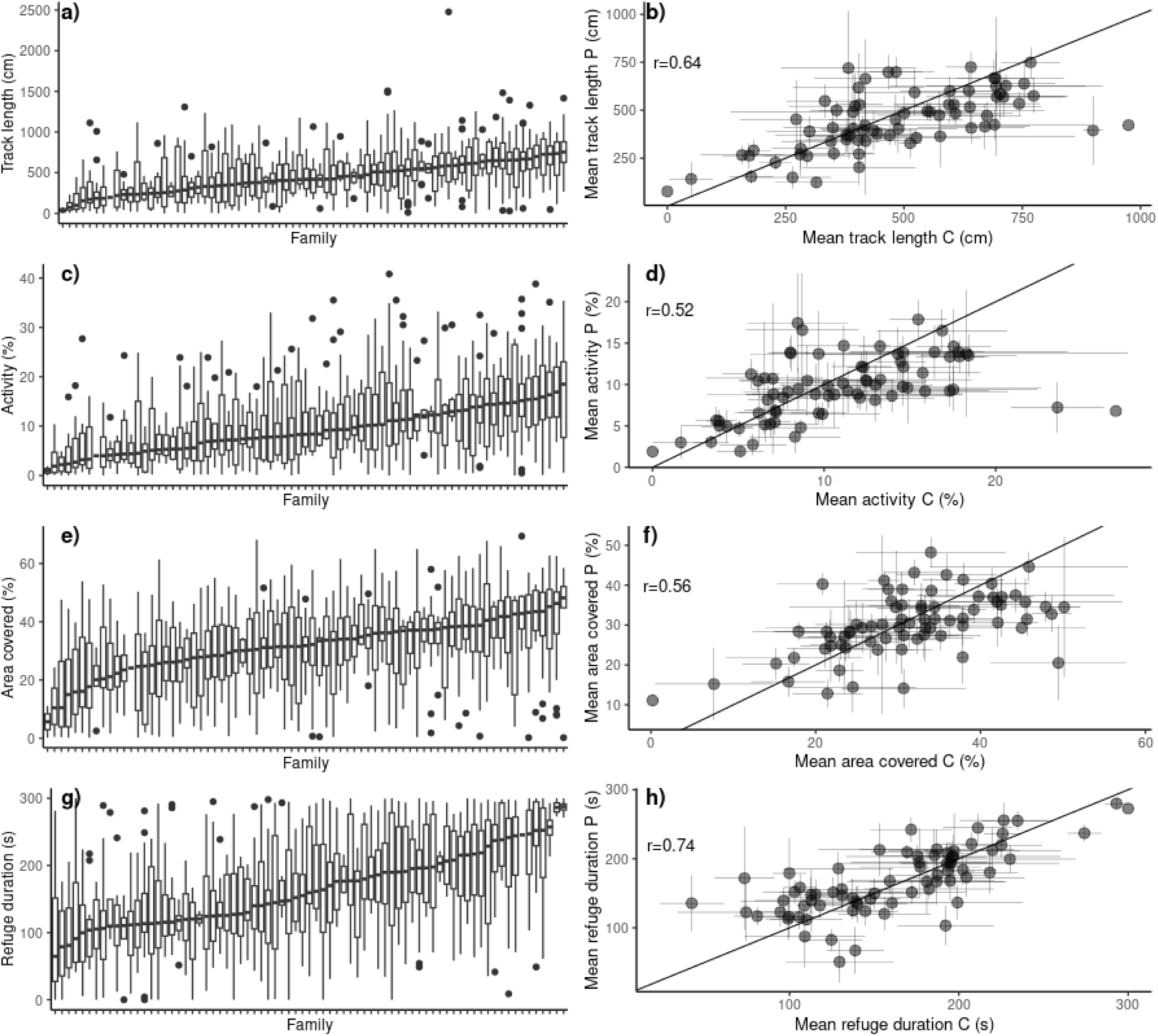
Behavioural variation among families. Shown are boxplots of each trait by family ordered by ascending trait median with both treatments combined (a,c,e,g). Also shown are pairwise plots of family mean behaviours by treatment (b,d,f,h) illustrating strong positive correlations. Vertical and horizontal error bars denote standard errors, while the diagonal 1:1 line is provided for reference.

**Table 3:** Univariate model comparisons for quantitative genetic data showing AIC and log-likelihood (LnL) for each model fitted to each trait as well as likelihood ratio tests (LRT) of among-family variance (V_F_), additive genetic variance (V_A_) and GxE. Shading denotes preferred model (lowest AIC) for each trait.

| Trait | Model | AIC | LnL | LRT Comparisons |  |  |  |  |
| --- | --- | --- | --- | --- | --- | --- | --- | --- |
| | | | | Comparison | Tests | $\chi^2$ | DF | P |
| TL | 2A | 1167.7 | -582.8 |  |  |  |  |  |
| | 2B | 1010.9 | -503.5 | 2.2 vs 2.1 | $V_F$ | 159 | 0,1 | <0.001 |
| | 2C | 1009.2 | -501.6 | 2.3 vs 2.2 | $V_A$ | 3.74 | 0,1 | 0.027 |
|  | 2D | 1013.2 | -501.6 | 2.4 vs 2.3 | GxE | 0.019 | 2 | 0.990 |
| AC | 2A | 1161.9 | -579.9 |  |  |  |  |  |
| | 2B | 1029.9 | -513.0 | 2.2 vs 2.1 | $V_F$ | 134 | 0,1 | <0.001 |
| | 2C | 1029.5 | -511.7 | 2.3 vs 2.2 | $V_A$ | 2.42 | 0,1 | 0.060 |
|  | 2D | 1033.5 | -511.7 | 2.4 vs 2.3 | GxE | 0.011 | 2 | 0.994 |
| AR | 2A | 1170.8 | -584.4 |  |  |  |  |  |
| | 2B | 1070.1 | -533.0 | 2.2 vs 2.1 | $V_F$ | 103 | 0,1 | <0.001 |
| | 2C | 1070.8 | -532.4 | 2.3 vs 2.2 | $V_A$ | 1.30 | 0,1 | 0.127 |
|  | 2D | 1072.7 | -531.4 | 2.4 vs 2.3 | GxE | 2.05 | 2 | 0.359 |
| RD | 2A | 1144.8 | -571.4 |  |  |  |  |  |
| | 2B | 972.2 | -484.1 | 2.2 vs 2.1 | $V_F$ | 175 | 0,1 | <0.001 |
| | 2C | 971.7 | -482.8 | 2.3 vs 2.2 | $V_A$ | 2.54 | 0,1 | 0.055 |
|  | 2.4 | 975.7 | -482.8 | 2.4 vs 2.3 | GxE | <0.001 | 2 | 1.000 |

Given the lack of support for GxE, we focus on genetic parameter estimates from the multivariate formulation of model 2C (Table 4). Though **G** is characterised by high uncertainty, point estimates of heritability (SE) calculated from this model ranged from h^2^ = 0.235 (0.224) for AR, to h^2^ = 0.281 (0.199) for RD and were much greater than the intra-class correlations associated with family (ICC_F_, interpretable as the proportion of variance explained by non-genetic, but family level, effects; Table 4). For the multivariate phenotype overall, additive genetic effects explained an estimated 24.3% of the total phenotypic variance conditional on fixed effects, while non-genetic family effects explained and estimated 7.0%. This indicates most the among-family variation visible in the raw data is attributable to genetic factors rather that common environment effects. Correlation structure in **G** recapitulates that in **ID;** r_G_ is strongly positive among TL, AC and AR, and strongly negative between these traits and TR. ***g_max_*** accounts for 96% of the variance in **G** (95% CI: 80-100%) and loads on the four original traits very similarly to ***id_max_*** (Figure 2, panels b vs c). The directional similarity of ***id_max_*** and ***g_max_*** is reflected in a low divergence angle (θ) of 14° and a vector correlation of 0.996.

**Table 4:** Additive genetic variance-covariance matrix **G** estimated from the quantitative genetic data set with trait-specific V_A_ on the diagonal (black shading), genetic covariance below the diagonal, and the corresponding genetic correlations (r_G_) above the diagonal (grey shading). Also shown are heritability estimates (h^2^) and the corresponding intra-class correlations for non-genetic family effects (ICC_F_). All estimates are from a multivariate formulation of model 2.3 which assumes an absence of GxE so that **G**, h^2^ and ICC_F_ are homogenous across treatments. Standard errors are shown in parentheses.

| Trait | **G** matrix | | | | $h^2$ | $ICC_F$ |
| --- | --- | --- | --- | --- | --- | --- |
|  | TL | AC | AR | RD |  |  |
| TL | 0.275 (0.184) | 0.959 (0.064) | 0.821(0.227) | -0.867 (0.206) | 0.279 (0.180) | 0.064 (0.080) |
| AC | 0.250 (0.174) | 0.232 (0.170) | 0.821 (0.213) | -0.707 (0.366) | 0.236 (0.168) | 0.064 (0.075) |
| AR | 0.208 (0.157) | 0.193 (0.152) | 0.179 (0.152) | -0.791 (0.280) | 0.235 (0.224) | 0.090 (0.082) |
| RD | -0.261 (0.181) | -0.233 (0.167) | -0.201 (0.159) | 0.274 (0.201) | 0.281 (0.199) | 0.084 (0.090) |

## Discussion

Here we investigated the quantitative genetics of personality and plasticity in a captive population of the red cherry shrimp, *Neocaridina davidi (syn. N. heteropoda).* Our results confirm the previously reported presence of substantial among-individual variation in shy-bold type behaviours hypothesised to influence predation risk (Rickward et al., 2024) and indicate an important genetic contribution to this. However, while we do find support for a plastic response to novel predator cues, we find no evidence that plasticity varies among individuals (IxE) or genotypes (GxE) in the population.

Analysis of the individual repeated measures data set provided evidence of behavioural plasticity in cherry shrimp. On average, the addition of predator cues causes a multivariate plastic response towards a behavioural phenotype we view as being ‘shyer’. This direction was expected, given that risk-taking behaviour is plastically moderated under elevated predation risk in many animals (see examples; Kim et al.,2016; Johnson & Sih, 2007; Sih et al., 2003). In decapods, directly comparable studies remain limited, but plasticity has been reported for behaviours functionally related to predator avoidance and/or escape. These include catatonic posture in crabs (Hazlett & Mclay, 2005), locomotory tail flips in crayfish (Bouwma & Hazlett, 2001) and startle response duration in hermit crabs (Briffa et al., 2008). Although little is known about the ecology of *Neocaridinia* in the wild, shrimp populations are generally subject to fish predation (Minello & Zimmerman, 1991; Salini et al., 1990) and risk aversion in the presence of fish cues seems likely to be an adaptive plastic response (Lima & Dill, 1990). Assuming so, it is perhaps surprising that the average effect size of predator cues is so modest (with a mean change of 0.237 SDU across the individual traits). Speculatively, the small effect size could reflect our methodology if, for example, the signal strength was low relative to concentrations of predator cues found in the wild. It may also be that stronger plastic responses would be observed if multiple cue types were present as many aquatic animals rely on multi-modal sensory detection of predators (Mukherjee et al. 2024). It is also possible that the limited response magnitude reflects costs of behavioural plasticity (e.g., lost resource acquisition opportunity from hiding) that are high relative to any expected benefit from reduced predation risk (Dall et al., 2004; Hazlett, 1995). Finally, noting *Xiphophorus helleri* represents a novel predator here, cherry shrimp lack eco-evolutionary experience with this species and so may be exhibiting sub-optimal, and potentially inadequate, plasticity as a consequence of prey naiveté (Carthey & Blumstein, 2018). If so, it is possible stronger plastic responses would be seen when presented with cues from a species of fish that represents a natural predator in the ancestral Taiwanese habitat.

While the repeated measures data corroborates our previous finding of among-individual differences in shy-bold behaviours (Rickward et al., 2024), analysis of the quantitative genetic dataset suggests among-individual differences arise, at least in part, from genetic factors. Thus, we see among-family variation consistent with genetic variance, but also find that family means are broadly similar, and strongly correlated, across treatments. Using the animal model, the vast majority of this among-family variation was attributed to additive genetic effects. Specifically, we estimated heritability of the multivariate behavioural phenotype to be 24% as compared to a family-level (non-genetic) intra-class correlation of just 7%. Our estimated heritability is moderately high for a behavioural phenotype, but well within the range of estimates reported in other studies (e.g., Brommer et al., 2012; Dingemanse et al., 2002; Kralj-Fišer & Schneider, 2012; Prentice et al., 2020). On average, additive genetic effects account for approximately half the personality variation (i.e. among-individual variance) found in animal populations(Dochtermann et al., 2019) though here it may be higher. From the repeated measures data we estimated that individual identity explains approximately 29% of the phenotypic variance in multi-trait space, while additive genetic effects explain 24% in the quantitative genetic data set. Accepting these as directly comparable implies that 82% of personality variation is attributable to genes (0.24/0.29 = 0.82).

As a partial caveat to our interpretation of genetic parameter estimates, we acknowledge that statistical uncertainty is high. At the outset, we had expected paternal half-sibships to provide more power to partition additive genetic from any non-genetic (full-sib) family effects present. However, despite breeding groups containing four females to each male, on average only two females per breeding group contributed to the offspring generation. To our knowledge, this study represents the first use of *Neocaridina* shrimp in a quantitative genetic breeding design and future studies might usefully increase the number of potential dams per sire to improve this. Our subsequent reliance on full-sib families housed in separate tanks also creates some risk that common environment (or maternal effects) may upwardly bias V_A_ (Kruuk & Hadfield, 2007). Thus, as well as increasing paternal half-sibship structure full siblings would ideally be mixed across tanks (e.g. following Houslay et al., 2022). However, doing this while retaining pedigree information requires permanently tagging individuals for identification and we have not yet found a suitable method for this. Nonetheless, within this constraint we did control for common environment effects as far as possible experimentally (e.g., by use of a recirculating system to standardise water) and statistically (e.g., by including a family-level random effect).

In general, prey species should be able to adapt to novel predators given sufficient time and heritable variation (Carthey & Blumstein, 2018). Here we find behaviour is repeatable and heritable in cherry shrimp such that, if novel predators do impose directional selection on shy-bold phenotype, we would predict adaptation of mean phenotype. However, we found no support for variation in the plastic response to predator cues among individuals (IxE) or among genotypes (GxE). This implies that the contribution of plasticity to any future adaptation will be limited. Specifically, in the absence of IxE, plastic responses are uniform across individuals and therefore cannot be selected on directly (i.e. viewing plasticity itself as a as a trait; Dall et al., 2019) or indirectly (via environment-specific selection on the trait displaying plasticity; Via & Lande, 1985). Moreover, even if IxE were present any further adaptation of plasticity would be contingent on the presence of genetic variation (GxE).

The absence of IxE in behavioural response to predator cues was somewhat surprising us as this phenomenon has now been widely demonstrated in empirical studies of shy-bold behaviours (e.g. Araya-Ajoy & Dingemanse, 2017; Houslay et al., 2018; Maskrey et al., 2018). However, given this result, the absence of GxE follows inevitably. Among-individual variance sets an upper limit on additive genetic variance (Falconer & Mackay, 1996) for plasticity just as it does for traits themselves. Previous tests of GxE in behaviours specifically linked to shy-bold personality remain rare, making it somewhat difficult to place the current result in a wider empirical context. Martin et al., 2017 demonstrated that genetic factors influence residual (i.e. putatively environmental) variance in behaviour of wild marmots at capture. (Prentice et al., 2023) reported a similar result for stress-related behaviour in guppies subject to open field trials. By demonstrating genetic variation in the degree of environmentally induced phenotypic variance, both of these actually show GxE on shy-bold traits (i.e. behavioural response to perceived risk). However, they do it via an approach that neither defines, nor tests, any specific environmental factor (i.e. the E remains unknown, see Prentice et al., 2023 for further discussion). More broadly however, there is support for GxE on labile animal behaviours across a range of environmental factors tested (e.g. Callander et al., 2013; Kasumovic et al., 2012; Rapkin et al., 2017). It would therefore be interesting to determine whether the absence of plasticity variation in cherry shrimp holds true across a wider set of behaviours (e.g. willingness to feed) and/or environmental factors likely to influence the relative costs and benefits of risk-taking behaviours. The latter might usefully include addition extrinsic cues of risk (e.g. visual presentation of predator models) but also intrinsic states (e.g. hunger level) likely to mediate risk-taking.

While many studies of personality variation rely on using single behavioural traits (e.g., time to emerge from a shelter to measure boldness or risk taking) two points emerge from our multivariate approach. First, multiple traits provide more information about individuals and the population level structure of behavioural variation. Arguably, those gains are limited by the strong correlation structure in **ID** and **G** in the sense that biological conclusions would have been very similar had they been based on just one trait (e.g. refuge duration). However, correlations were not known *a priori*. Rather they were expected, subject to the implicit assumptions that (i) there is an axis of shy-bold variation among individuals in this species and (ii) the chosen traits would reflect this axis as they have been shown to do in other systems. Thus, the estimated correlation structures actually provide useful corroboration of assumptions about a system we currently know little about. It is equally true that failure to corroborate starting expectations can sometimes provide valuable biological insights into the structure and putative function of behavioural variation (Houslay et al., 2022; Rickward et al., 2024). The second major advantage of our multivariate approach is that it allows us to compare the shape of behavioural variation across multiple levels. Specifically, we found close directional alignment between the vector of multivariate plasticity (i.e. the average change in multivariate behaviour when exposed to predator cues), and both **id_max_** and **g_max_** - the main axes of among-individual and additive genetic variance respectively. Alignment of plasticity with ***id_max_*** provides further ‘face validity’ (sensu Burns, 2008) for our assay. This is because shy-bold variation is defined as behavioural response to risk, while our predator treatment was intended to increase the perception of that risk. While our study was not intended as a direct test of either hypothesis, we also note that the alignment of plasticity with **g_max_** is predicted if plasticity is adaptive (Berdal & Dochtermann, 2019), and that the similarity of **ID** and **G** supports Cheverud’s conjecture that phenotypic correlations will typically be a good proxy for genetic correlations (Cheverud, 1988; Dochtermann, 2011)

In summary, we find support for genetic variance underpinning among-individual differences in shy-bold behaviour in cherry shrimp, *Neocaridina davidi*. We also find evidence of plasticity, with individuals shifting towards a ‘shyer’ most risk averse average phenotype in the presence of cues from a novel (fish) predator. However, we found no variation among individuals or genotypes in their plastic responses. This implies that average shy-bold behaviour should evolve if subject to selection by predators as is widely hypothesised. Nevertheless, our results suggest the further adaptive evolution of behavioural plasticity may be constrained. Testing these predictions using experimental evolution approaches should be feasible given the ease of husbandry, small size, and short generation time of this emerging decapod model. This would allow rare insight into the evolutionary dynamics of animal personality under experimentally manipulated selection regimes. At the same time, very little is known about how predation shapes selection on shrimp behaviour in the wild -either in their native range or areas where invasive populations have established. Does natural selection generally favour shy or bold phenotypes? Does this depend on predation risk? Could predation mediate a risk-reward trade-off that maintains variation? Field-based studies to address these gaps in our knowledge would provide valuable ecological context for our results.

## Acknowledgements

We thank Tom Black and the animal care staff in the aquatic facility for assistance with shrimp husbandry. FS was supported by a MSCA postdoctoral fellowship.

## Ethics statement

All data for this study were collected under local ethical approval (University of Exeter approval ID 517031)

## Supplemental Appendix: Character state modelling of GxE

In the main text we present tests of GxE formulated using random regression animal models (model 2.4) that characterise genetic ‘reaction norms’ with two parameters – an intercept (genetic merit in control conditions), and a slope on the ‘environment’ of treatment (arbitrarily scaled numerically).

The latter is interpreted as genetic merit for plasticity. Genetic variance in slope (plasticity) is GxE. An alternative ‘character state’ approach is to consider treatment-specific behaviours as potentially distinct phenotypes. Though formally equivalent, reaction norm and character state frameworks can provide complementary perspectives and insights (see e.g. Roff and Wilson 2014 for a didactic treatment).

### Supplemental character state models fitted

Under the character state view, for a single behaviour, GxE is manifest as a change in V_A_ across environments and/or a cross-environment genetic correlation <+1. We therefore applied the character state perspective to the quantitative genetic dataset, by defining two environment-specific characters for each of our original behavioural traits. Each pair of environment-specific characters (e.g. TL*_(c)_* and TL*_(p)_*) was then analysed with a bivariate animal model containing fixed effects as described for Models 2A-2D, and random effects of additive genetic and family-level intercepts.

Genetic and family covariance were estimated as 2×2 unstructured matrices containing treatment specific variances and the cross-treatment covariance. We rescaled the genetic covariance to the cross-treatment genetic correlation for ease of interpretation. Residuals were modelled with a diagonal matrix containing treatment specific residual variances but no covariance. This is because residual (observation level) covariance is not identifiable when the two response variables are not simultaneously observed.

For a multivariate behavioural phenotype, GxE could equally be evidenced by treatment-sensitivity of the genetic variance-covariance matrix such that **G***_(c)_* ≠ **G***_(p)_.* Given high uncertainty on all genetic parameter estimates it is clear that any statistical comparison of **G***_(c)_* and **G***_(p)_* estimated from the currently available data would be very underpowered. Nonetheless, we fitted a four-trait animal model to each set of treatment-specific characters (with fixed effects as used in all models) to obtain point estimates of **G***_(c)_* and **G***_(p)_* for qualitative comparison. Note that in principle it is possible to jointly model all 8 characters at once (i.e. 4 behaviours x 2 treatments), to jointly estimate treatment specific **G** matrices, and all cross-treatment genetic covariances (within- and among-behaviours). In practice we elected not to do this as we felt data volume was sure to be limiting.

### Results and conclusions arising

Bivariate animal model further supported the conclusion that GxE are absent, or at least of limited magnitude. Though we did conduct formal inference on GxE, point estimates of V_A(*c)*_ and V_A*(p)*_ are very similar for each trait (Table S2) and approximate 95% CI (calculated as parameter estimates ±1.96*SE) are very strongly overlapping. Similarly, point estimates of the cross-context genetic correlation are close to unity in each case (with approximate 95% CI bracketing +1). While reiterating that no statistical comparison is presented, the two 4 trait models yielded REML point estimates of **G***_(c)_* and **G***_(p)_* that are extremely similar (Table S1). These estimates have a matrix correlation of r=+0.993, while the first eigen vectors of these two matrices have a vector correlation of 0.999 and a divergence angle (θ) of 3.4°. Thus, the two estimates are qualitatively indicative of an absence of GxE.

**Table S1-.** Estimated fixed effects with statistical inference from univariate models of each trait. Estimates are from Model 1.3 fitted to the repeated measures data set (see main text)

| Trait | Effect (Level) | Coefficient (SE) | F | DF | P |
| --- | --- | --- | --- | --- | --- |
| TL | *intercept* | 1.262 (1.788) | 0.175 | 1, 76.6 | 0.677 |
|  | *treatment (P)* | -0.252 (0.092) | 7.547 | 1, 395.1 | 0.006 |
|  | *temperature* | -0.006 (0.078) | 0.006 | 1, 409.3 | 0.938 |
|  | *order* | -0.001 (0.015) | 0.010 | 1, 413.0 | 0.921 |
|  | *arena (2)* | -0.068 (0.117) | 0.713 | 3, 415.4 | 0.544 |
|  | *(3)* | 0.059 (0.143) |  |  |  |
|  | *(4)* | 0.105 (0.152) |  |  |  |
|  | *age<sub>family</sub>* | 0.002 (0.002) | 0.614 | 1, 77.6 | 0.436 |
|  | *n<sub>family</sub>* | -0.019 (0.008) | 5.593 | 1, 77.3 | 0.021 |
|  | *repeat* | -0.101 (0.023) | 18.44 | 1, 443.2 | <0.001 |
|  | *time* | -0.001 (0.001) | 2.420 | 1, 409.4 | 0.121 |
| AC | *intercept* | 1.140 (1.762) | 0.240 | 1, 76.5 | 0.626 |
|  | *treatment (P)* | -0.257 (0.090) | 8.114 | 1, 394.6 | 0.005 |
|  | *temperature* | 0.002 (0.077) | 0.001 | 1, 408.0 | 0.974 |
|  | *order* | -0.001 (0.014) | 0.004 | 1, 411.5 | 0.952 |
|  | *arena (2)* | -0.139 (0.116) | 1.202 | 3, 413.7 | 0.309 |
|  | *(3)* | -0.105 (0.141) |  |  |  |
|  | *(4)* | 0.033 (0.149) |  |  |  |
|  | *age<sub>family</sub>* | 0.002 (0.002) | 0.868 | 1, 77.4 | 0.355 |
|  | *n<sub>family</sub>* | -0.022 (0.008) | 6.774 | 1, 77.1 | 0.011 |
|  | *repeat* | -0.101 (0.023) | 19.14 | 1, 445.4 | <0.001 |
|  | *time* | -0.001 (0.001) | 2.574 | 1, 408.1 | 0.109 |
| AR | *intercept* | -0.956 (1.817) | 0.219 | 1, 76.4 | 0.641 |
|  | *treatment (P)* | -0.059 (0.093) | 0.398 | 1, 394.9 | 0.529 |
|  | *temperature* | 0.081 (0.079) | 1.049 | 1, 409.1 | 0.306 |
|  | *order* | 0.013 (0.015) | 0.722 | 1, 412.8 | 0.396 |
|  | *arena (2)* | -0.100 (0.119) | 1.528 | 3, 415.2 | 0.207 |
|  | *(3)* | -0.014 (0.146) |  |  |  |
|  | *(4)* | 0.149 (0.154) |  |  |  |
|  | *age<sub>family</sub>* | 0.003 (0.002) | 2.162 | 1, 77.4 | 0.146 |
|  | *n<sub>family</sub>* | -0.019 (0.008) | 5.350 | 1, 77.1 | 0.023 |
|  | *repeat* | -0.086 (0.024) | 12.90 | 1, 443.2 | <0.001 |
|  | *time* | -0.001 (0.001) | 4.451 | 1, 409.2 | 0.035 |
| RD | *intercept* | -3.64 (1.919) | 4.024 | 1, 76.0 | 0.048 |
|  | *treatment (P)* | 0.380 (0.100) | 14.55 | 1, 399.9 | <0.001 |
|  | *temperature* | 0.075 (0.084) | 0.785 | 1, 422.7 | 0.376 |
|  | *order* | -0.026 (0.016) | 2.647 | 1, 428.3 | 0.104 |
|  | *arena (2)* | 0.224 (0.126) | 2.981 | 3, 431.8 | 0.031 |
|  | *(3)* | -0.028 (0.154) |  |  |  |
|  | *(4)* | -0.134 (0.163) |  |  |  |
|  | *age<sub>family</sub>* | 0.001 (0.002) | 0.117 | 1, 77.4 | 0.734 |
|  | *n<sub>family</sub>* | 0.016 (0.007) | 5.536 | 1, 77.0 | 0.021 |
|  | *repeat* | 0.055 (0.025) | 4.849 | 1, 425.0 | 0.028 |
|  | *time* | 0.002 (0.001) | 11.58 | 1, 423.2 | 0.001 |

**Table S2:** Estimated treatment-specific variances (V_A_) where subscripts *(c)* and *(p)* denote control and predator treatments. Also shown is the estimated cross-treatment genetic correlation for each behaviour (r_G*(cp)*_). Standard errors are shown in parentheses.

| Trait | $V_{A(c)}$ | $V_{A(p)}$ | $r_{G(cp)}$ |
| --- | --- | --- | --- |
| TL | 0.274 (0.217) | 0.254 (0.199) | 0.946 (0.135) |
| AC | 0.244 (0.224) | 0.207 (0.175) | 0.947 (0.161) |
| AR | 0.166 (0.191) | 0.144 (0.167) | 0.855 (0.272) |
| RD | 0.340 (0.238) | 0.298 (0.253) | 0.952 (0.238) |

**Table S3:** Point estimates of treatment-specific **G** matrices under control *(c)* and predator *(p)* treatments showing additive genetic variances on the diagonal (black shading), covariances below the diagonal, and genetic correlations above the diagonal (grey shading).

| Trait | $\mathbf{G}_{(c)}$ | | | | $\mathbf{G}_{(p)}$ | | | |
| --- | --- | --- | --- | --- | --- | --- | --- | --- |
|  | TL | AC | AR | RD | TL | AC | AR | RD |
| TL | 0.361 | 0.994 | 0.963 | -0.977 | 0.254 | 0.982 | 0.990 | -0.891 |
| AC | 0.326 | 0.297 | 0.970 | -0.965 | 0.237 | 0.229 | 0.998 | -0.831 |
| AR | 0.279 | 0.255 | 0.233 | -0.945 | 0.176 | 0.168 | 0.124 | -0.854 |
| RD | -0.351 | -0.315 | -0.273 | 0.358 | -0.247 | -0.219 | -0.166 | 0.303 |

